# Recreating coronary vascularization and sympathetic innervation of myocardium on a human pluripotent stem cell-derived heart organoid

**DOI:** 10.1101/2025.01.10.632325

**Authors:** Mariana A. Branco, Mafalda Marques Nunes, Ana Luísa Rayagra, Miguel F. Tenreiro, Joaquim M.S. Cabral, Maria Margarida Diogo

## Abstract

Coronary vascularization and sympathetic innervation of the myocardium is a concomitant event during embryonic heart development and both systems are crucial to ensure normal adult heart function. Here we describe a self-organized hiPSC-derived heart organoid that recreates both the coronary vascular plexus and the sympathetic neuronal network of the ventricle myocardium, with a physiologically relevant in-vivo-like structural organization and function. Through modulation of PDGF-β and VEGF signalling pathways, we attained a heart organoid that incorporates 1) an external epicardial layer (mesothelium) of DACH1, NR2F2 and WT1 positive cells, 2) a sub-epicardial space from where a functional primary coronary vascular plexus of CD31^+^/DACH1^+^ cells emerge, 3) a compact myocardial region adjacent to the epicardium, enriched in proliferative cardiomyocytes and ECM deposition, and 4) a sympathetic neuronal network that controls heart organoid contraction. Therefore, the human heart organoid described herein, is a unique model to study new regenerative medicine-based approaches to restore innervation and promote re-vascularization in adult heart after ischemic events and to perform adult and developmental cardiotoxicity studies.

## INTRODUCTION

Coronary vascularization and innervation of the myocardium is an interlinked event and a crucial stage in embryonic heart development. Healthy coronary vasculature and sympathetic innervation of the myocardium are crucial to ensure normal heart function. Sympathetic signals control heartbeat rate, relaxation time, contractile force, and conduction velocity, and the coronary vascularization ensures the proper oxygen demand to the working myocardium. Dysfunction in one or both cell systems is the most prevalent cause of heart failure, arrhythmogenesis, and sudden cardiac death. Therefore, the development of physiologically relevant human induced pluripotent stem cell (hiPSC)-derived heart organoid models incorporating both coronary vascularization and sympathetic innervation provides an opportunity to dissect the mechanisms and find new therapies for heart pathologies in a clinically relevant context.

The in vitro recreation of the combined developmental process of heart coronary vascularization and sympathetic innervation in hPSC-derived heart organoid models has not been achieved yet. Although there are several studies describing hPSC-derived vascularized cardiac models ^1–4^, none recreates the spatiotemporal process of the early coronary vascular plexus development, nor its interaction with the sympathetic ganglionated network. Also, the establishment of hPSC-derived heart organoids recreating only myocardium innervation has not been addressed yet, with only a few studies exploring co-culture strategies between primary or hPSC-derived ventricle cardiomyocytes (CMs) and sympathetic neurons (SNs) in 2D monolayer systems ^5–14^. Therefore, there is a lack of physiologically relevant in vitro models that recreate both coronary vascularization and innervation.

In the present work, we follow an in-vivo inspired embryonic developmental approach and take advantage of our previously established epicardium-myocardium organoid (EMO) ^15^ as the starting model to develop a stepwise platform that recreates the development of heart coronary vascularization in syntony with SNs innervation. The developed heart organoid model 1) allows unprecedented access to the mechanisms involved in the establishment, function and regeneration of human ventricle myocardium coronary vasculature and innervation, and 2) can be explored in clinically needed areas, specifically to assess adult and developmental cardiotoxicity, to disclose new therapeutic strategies to target arrhythmogenic cardiomyopathies through autonomic nervous system modulation, and to develop new regenerative medicine-based approaches to restore innervation and promote re-vascularization in adult heart after ischemic events.

## RESULTS

Previous work developed by our group has reported an hiPSC-derived EMO model (control EMOs), featuring a self-organized epicardium region surrounding the entire surface of a ventricle myocardium-like area^15^ (**Figure S1A-S1D**). Knowing that throughout embryonic heart development, epicardium formation proceeds with the coronary vascularization and innervation of the myocardium, we hypothesized that EMOs could be a scaffold to explore the development of a coronary vascularized and innervated heart organoid.

### Vascular network in EMOs mimics the primitive coronary vascular plexus in the developing human heart

To induce vascularization, we started by assessing EMO’s response to vascular endothelial growth factor (VEGF) and platelet-derived growth factor-β (PDGF-β) signalling pathways modulation (**Figure S2A**). PDGF-β signalling has been described to be required for efficient epicardium cell migration and derivation into coronary vascular smooth muscle cells in vivo ^16,17^ and it has been used to induce vascular smooth muscle cells specification from hiPSC-derived epicardial cells in vitro ^18^. On the other hand, VEGF is an angiogenic factor that is commonly used as an inducer of endothelial cells (ECs) specification and maintenance. Thus, the supplementation of PDGFBB and VEGFA was tested during the process of EMOs generation.

To perform the characterization of VEGFA- and PDGFBB-treated EMOs, we assessed the expression of DACH1 marker, which has been reported to specifically label coronary vessels in a mouse model ^19–22^. Also, knowing that the epicardium comprises a heterogeneous cell population of multipotent progenitors, combining mesothelial cells, which line the cardiac surface, and cells that exhibit a predisposition towards epithelial-mesenchymal transition (EMT) ^23^, we complemented the characterization of the epicardium and epicardium-derived cells within our model using additional markers apart from the commonly used WT1. Specifically, we assessed the expression of NR2F2 (COUP-TFII), which is commonly observed in mesenchymal cells during organogenesis, playing a regulatory role in angiogenesis and heart development, and it has been shown to be required for proper sinus venosus patterning, the main cell source of coronary ECs, via mesenchymal–endothelial signalling ^24,25^. Additionally, we also assessed the expression of 1) NG2, which identifies epicardium-derived mesenchyme region and pericytes ^26^, 2) the mesenchymal/cardiac fibroblast marker vimentin (VIM) and the 3) ECM proteins laminin (LAM) and fibronectin (FIB), to deeper characterize the extent of EMT of epicardial progenitor cells.

The supplementation of VEGFA during EMOs generation (V-EMOs) resulted in a significant increase of CD31^+^ ECs (from 1.6±0.5% to 3.9±0.2% of the total EMOs’ area) (**Figure S2B**), which form a network of connected tubules at the interface of epicardial region and myocardial area (**Figure 1A**, **Figure S2C, Video S1**), with branches sprouting toward the myocardium zone (**Figure 1B and Figure S2D**). We also observed that the EC-network is lined by vascular smooth muscle cells/pericytes (NG2^+^ cells) and the ECM protein laminin, essential to form a functional vascular plexus (**Figure 1C and Figure S2E**). Additionally, ECs present a positive nuclear staining for both WT1 and DACH1 markers and were negative for the venous marker NR2F2, suggesting that the vascular cells can be representative of the primordial stage of coronary vascular plexus (**Figure 1D and 1E, Figure S2F**). In V-EMOs, as in control EMOs, the myocardial region is covered in almost all the extent of the organoid surface by a thin layer of cells, with the presence of thicker regions, in smaller extents of the organoid surface, both layers comprising NR2F2, DACH1, and WT1 positive cells (**Figure S2G**).

**Figure 1:**
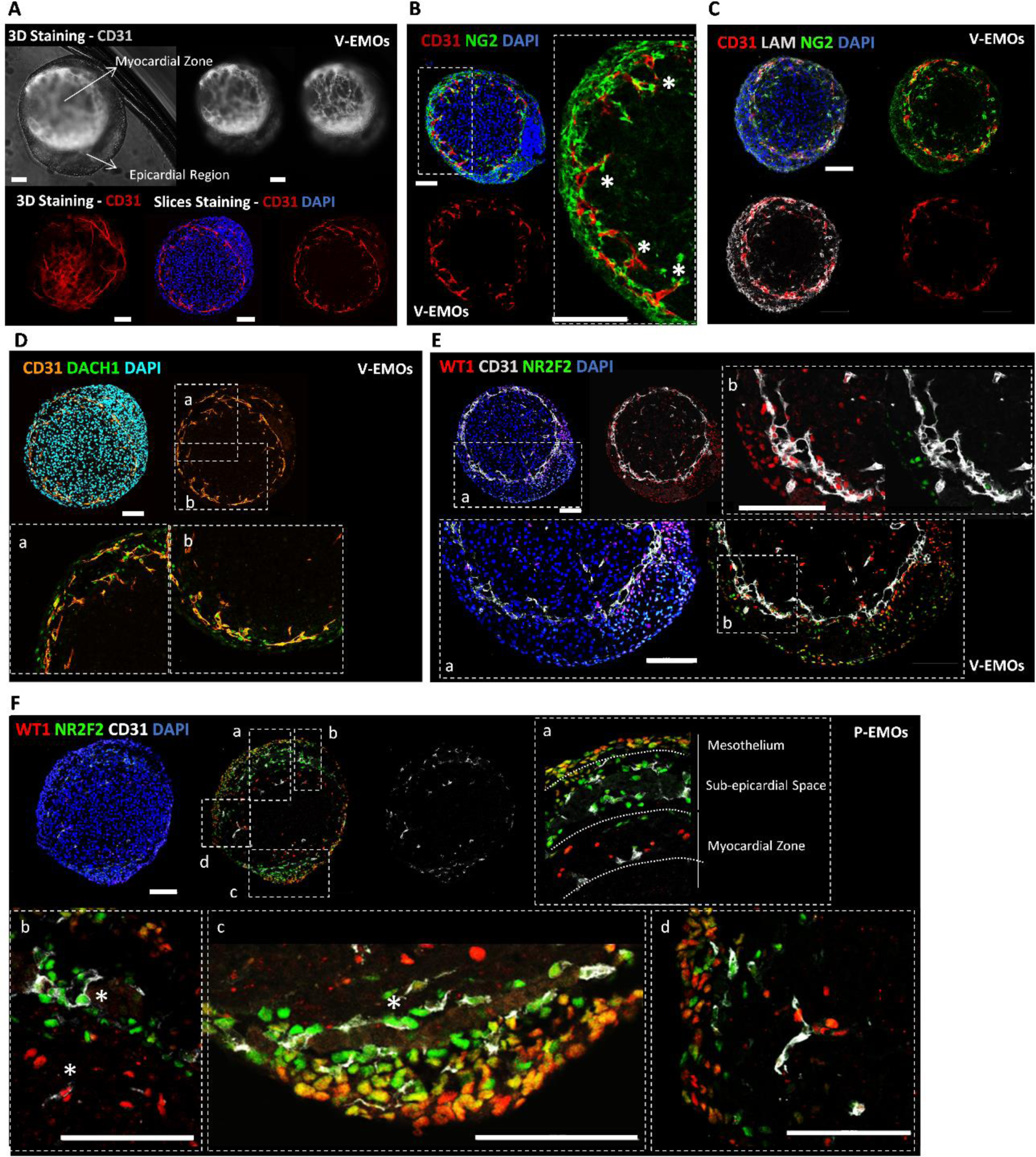
EMOs stimulation with VEGFA and PDGFBB induces the formation of a coronary-like vascular network. **(A)** Representative IF staining of V-EMOs (3D and organoid slices), highlighting the development of a CD31^+^ network of vascular cells surrounding the myocardium region. **(B-E)** Representative IF staining of V-EMOs (organoid slices), highlighting the sprouting of CD31^+^ cells towards the myocardium region (B), the co-staining of CD31 network with the ECM laminin and the pericyte marker NG2 (C), and the staining for DACH1, WT1 and NR2F2 in CD31^+^ cells (D and E), suggesting a coronary vascular-like phenotype for this population. **(F)** Representative IF staining of P-EMOs (organoid slices), highlighting the generation of two distinct layers in the epicardium region, the mesothelium (WT1^+^/NR2F2^+^/DACH1^+^ cells) and the sub-epicardium space (WT1^−/+low^, NR2F2^+^ and DACH1^+^ cells) (a) and the two distinct sub-populations of ECs, namely CD31^+^/NR2F2^+^ cells within the sub-epicardium space and in the myocardium surface, and CD31^+^/WT1^+^/NR2F2^−^ cells within the myocardium region (b-d). Scale bars, 100 µm. See also Figure S1, S2 and S3.

In PDGFBB treated EMOs (P-EMOs), we observed a thick epicardium region, which resulted in EMOs with an increased diameter (696±18 µm EMOs vs 762±35 µm P-EMOs) (**Figure S3A – S3C**), suggesting an effect of PDGFBB in promoting EMT of epicardial cells. The epicardium region shows two clear distinct areas (**Figure S3D**), namely 1) a WT1^+^/NR2F2^+^/DACH1^+^ mesothelium-like layer and 2) a sub-epicardial region, with a lower cell density, which comprises WT1^−/+low^, NR2F2^+^ and DACH1 cells (**Figure 1F and Figure S3E**). Although with a less complex network of tubules in comparison to V-EMOs, staining for CD31 showed the presence of ECs, mainly located within the sub-epicardial region and in the outermost myocardium region, adjacent to the epicardium (**Figure 1F, Figure S3E and S3F)**. Interestingly, and contrarily to what was observed in V-EMOs, the ECs located in the subepicardial space present a NR2F2^+^/WT1^−^/DACH1^+^ phenotype (**Figure 1F**). Similarly to V-EMOs, the organized ECs within the myocardium region present a NR2F2^−^/WT1^+^/DACH1^+^ phenotype (**Figure 1F**). In vivo, it is known that part of the endothelial progenitors from the initial vascular plexus that develops at the subepicardial space continues to spread and mature at the myocardium surface, forming the coronary veins (NR2F2^+^ [15]), and some invade the myocardium, forming a branching network of coronary arteries [19,20]. Although in V-EMOs most of the ECs are NR2F2^−^/WT1^+^, in P-EMOs, we observed CD31^+^/NR2F2^+^ within the subepicardial space and in the myocardial surface, and CD31^+^/WT1^+^/NR2F2^−^ cells within the myocardial region, suggesting that P-EMOs may be recapitulating both the formation of coronary veins that mature at the myocardial surface, and intramyocardial coronary arteries specification. It is also possible that the CD31^+^/NR2F2^+^ ECs observed at the myocardial surface are still in the process of losing the venous phenotype before specification into coronary arteries, as occurs in vivo ^27^.

Regarding the myocardium zone, similarly to what has been observed in control EMOs, we confirmed the presence of a compact-like myocardial region adjacent to the epicardial zone, which comprises predominantly proliferative CMs (NKX2.5^+^/Ki-67^+^), with a significant increase in the percentage of these CMs in the compact myocardium zone in V-EMOs compared with control (from 8.8±1.0% in EMOs to 19.7±2.0% in V-EMOs of NKX2.5^+^/Ki-67^+^within the myocardium region) (**Figure 2A and Figure S4A**). Additionally, although this was also observed in control and V-EMOs (**Figure S4A**), the stronger staining for NG2 and VIM in the compact region of the myocardium, compared to the inner core, was more evident in P-EMOs (**Figure 2B and 2C**). P-EMOs also stained for the ECM proteins fibronectin and laminin, within the compact myocardium region, and externally, covering the myocardium zone (**Figure 2C and 2D**). Both observations suggest a more active EMT of epicardial progenitor cells into fibroblasts and pericytes, and its migration towards the myocardium, in the PDGFBB-supplemented condition. Interestingly, it was within the compact myocardial region of P-EMOs that it was observed an enrichment in CD31^+^ ECs (**Figure 2D).**

**Figure 2:**
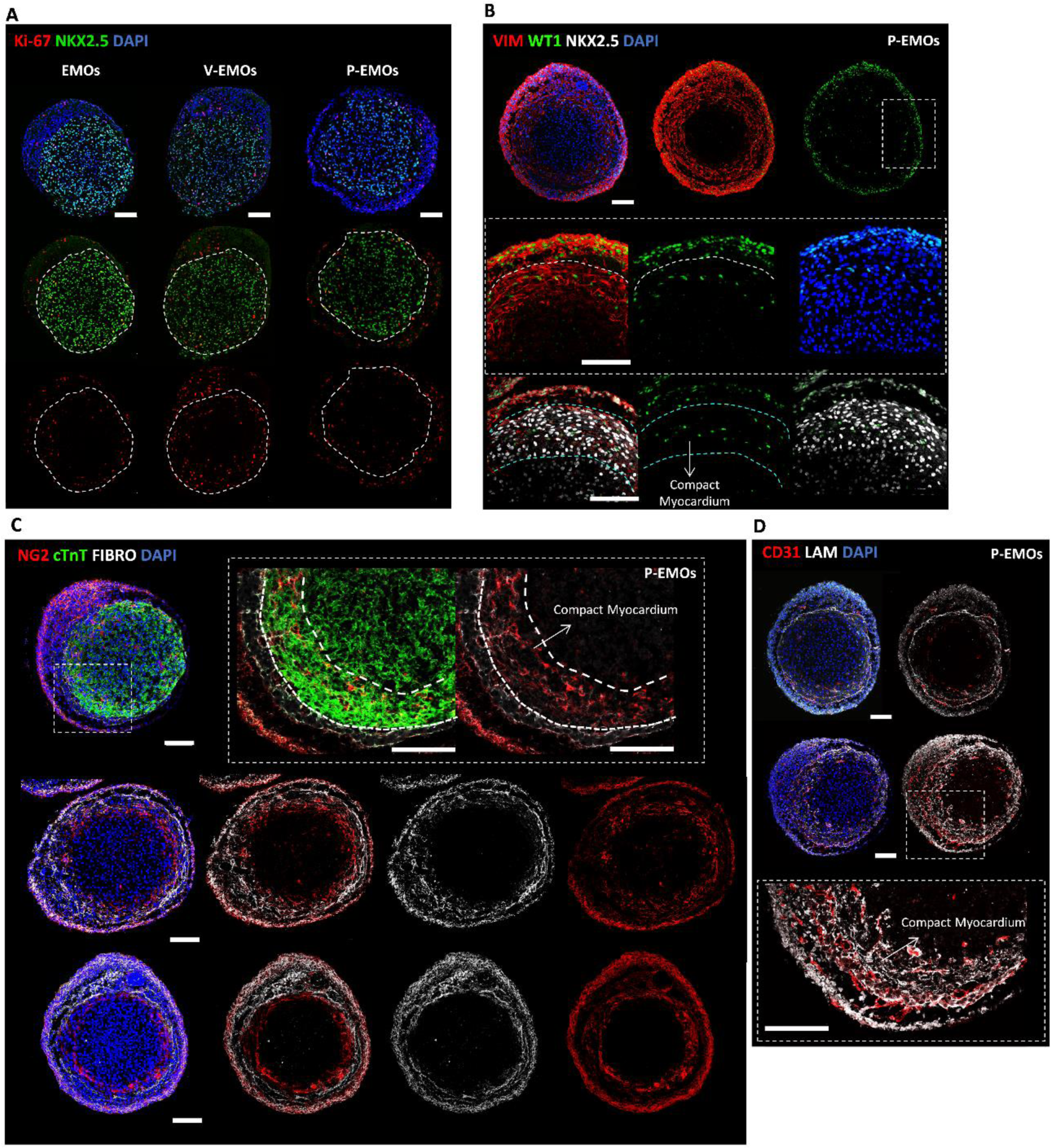
Improved vascularization and EMT of epicardium progenitors’ cells promote the development of a compact/proliferative myocardium region in EMOs. **(A)** Representative IF staining of control, V- and P-EMOs (organoid slices), highlighting the presence of proliferative CMs (NKX2.5^+^/Ki-67^+^), within the compact myocardium region. **(B-D)** Representative IF staining of P-EMOs, highlighting the staining of NG2 and VIM (B and C) and the presence of CD31^+^/WT1^+^ ECs (D), within the compact myocardium region. Scale bars, 100 µm. See also Figure S4.

### Assembly of V- and P-EMOs and SNS spheroids generates a functional heart-innervated organoid

To generate an innervated ventricle myocardium organoid model (iEMOs), we first established a differentiation platform to attain SNSs from hiPSCs. To achieve that, we adapted a previously established 2D differentiation protocol ^26^ to a 3D environment. Success of differentiation was confirmed by RT-qPCR analysis, that demonstrated a significant up-regulation of the SNs’ markers SOX10, ASCL1 and PHOX2B at the end of neural induction (Day 12 of differentiation) (**Figure S5A**) and by immunostaining for the SN enzyme tyrosine hydroxylase (TH) and the peripheral neuronal marker peripherin (PRPH) after 30-40 days of differentiation (**Figure 3A**). The functionality of SNs was assessed, using a MEA system, after 50 days of 3D differentiation followed by 8-12 additional days of culture on the electrodes. This analysis confirmed the electrical activity of the neurons (**Figure 3B and Figure S5B**), with these cells presenting an average neuron conduction velocity of 0.52±0.03m/s and a neuron firing rate of 3±0.3Hz. We then fused V- and P-EMOs with fluorescently-labelled SNSs (>D50 of differentiation) in 96-well plates to provide innervation to our heart organoid model. After 48 hours post-assembly, it was already possible to observe the beginning of neuronal projections towards the EMOs region of the assembloids (**Figure 3C**). Interestingly, after 6 and 12 days of assembly, a network of SNs densely formed at the surface of myocardial region (**Figure 3D and 3E**), below the epicardial zone. After 2 weeks of co-culture, we examined the effect of SN stimulation on contraction activity of EMOs to assess the presence of a functional interaction between ventricle CMs and SNs. To do so, we treated assembloids with nicotine, which activates nicotinic acetylcholine receptors of post-ganglionic neurons in the autonomic nervous system (ANS), and assessed beating rate of EMOs after 30 min with calcium imaging. We registered a significant increase in beating rate upon nicotine treatment, from 16±1 to 31±2 BPM (**Figure 3F**), confirming that our SNSs are capable of modulating cardiomyocyte function.

**Figure 3:**
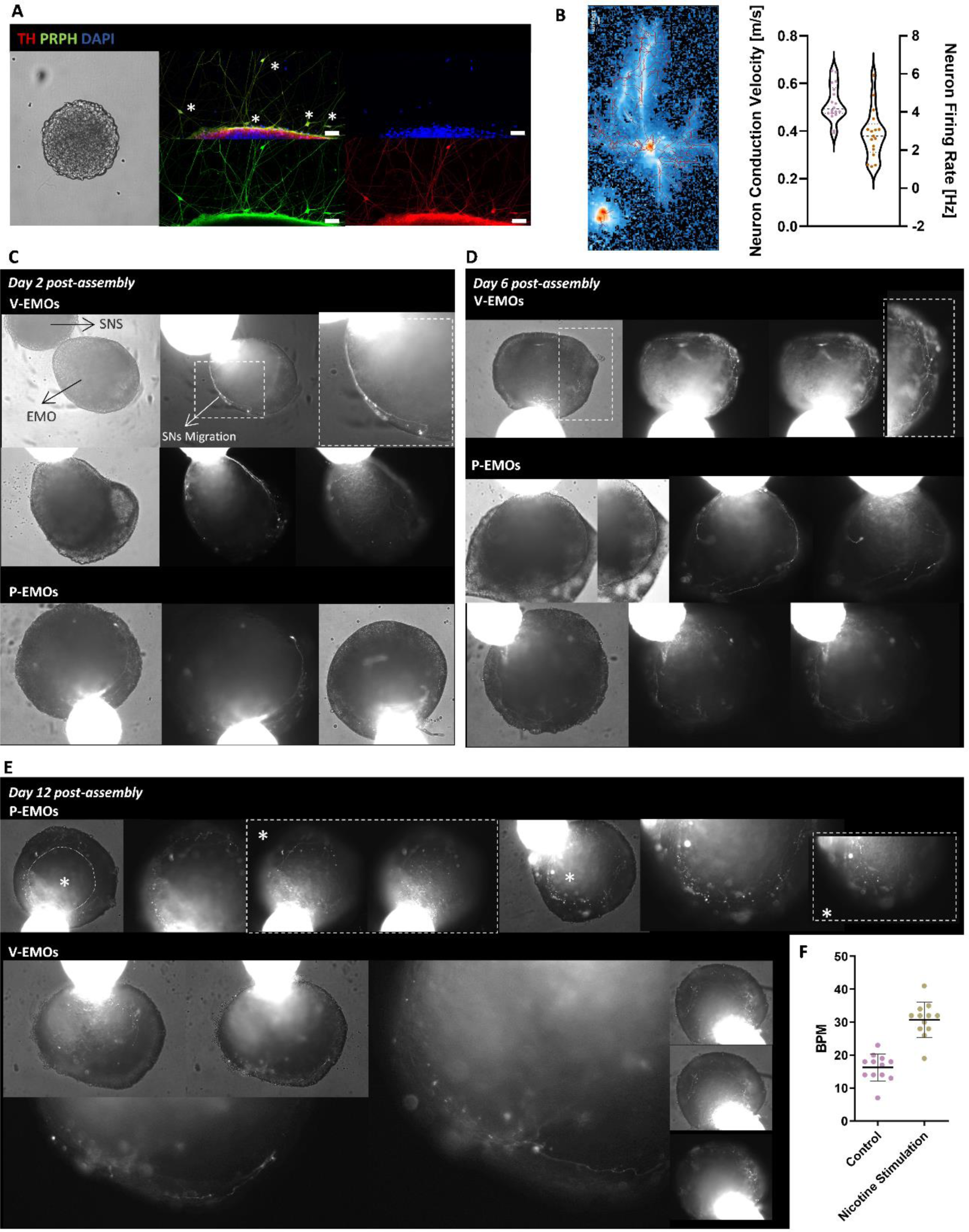
EMO-SNS assembloids show in-vivo-like functional sympathetic myocardium innervation. **(A)** Representative BF images of SNSs and IF staining of replated SNSs (7 days after replating) after 40 days of differentiation, highlighting the expression of sympathetic neuronal markers TH and PRPH. Scale bars, 100 µm. **(B)** Neuronal conduction velocity (m/s) and neuron firing rate (Hz) of SNSs, after 50 days of differentiation and 8-12 days of culture in MEA. Data are represented as mean ± SEM of n = 3 independent experiments. **(C-E)** Representative BF images of V- and P-EMO-SNS assembloids 2- (C), 6- (D) and 12-days (E) post-assembly, highlighting SNs migration from SNS region towards the EMOs region of EMOs. **(F)** Quantification of BPM of EMOs-SNS assembloids 10 days post-assembly, before and after drug stimulation with nicotine. Data are represented as mean ± SEM of n = 3 independent experiments. See also Figure S5.

## DISCUSSION

Normal heart function is ensured by a healthy myocardium supported by a functional vasculature and a functional autonomic nervous system. Alterations to the structure and function of the coronary vascular network and in the excitability, density, distribution, and neurotransmitter content of cardiac sympathetic fibers, are well described as events that compromise cardiac function. Therefore, the development of in vitro heart organoid models that recreate not only the myocardium but also the complementary cell systems that support the normal function of the heart is crucial to achieve an impactful model for different clinical and biomedical applications.

Herein, we describe the first hiPSC-derived physiologically relevant heart organoid that recreates both the coronary vascular plexus and the SNs network with an *in-vivo*-like structural organization and function. Although previously reported hiPSC-derived heart organoids have been shown to recapitulate at some extend the cardiac vasculature/endocardial vascularization ^28–31^, there is no model that accurately recapitulates coronary EC formation via the series of orchestrated events described to occur in vivo, being this a limitation identified in recent hiPSC-derived heart organoid models ^32^, nor recapitulates its interplay with the establishment of a functional innervated network. The process of development of both systems is still not completely disclosed, and most of the available information has been attained from studies performed in animal models. Despite that, it has been shown that the epicardium is an important source of paracrine factors and progenitor cells for the establishment of a functional coronary vasculature in the human heart ^26^. Also, it has been reported that sympathetic ventricle innervation develops in close cross-talk with vascularization, with neurotrophic factors secreted by epicardium-derived vascular cells guiding the extension of axons through the subepicardium and inducing invasion into the myocardium ^14,33^. In the present work, we used EMOs as the starting point to explore a developmentally inspired strategy to recreate the in-vivo like generation and interaction between epicardium, coronary vascular cells and SNs. Through exogenous manipulation of PDGF-β and VEGF signalling pathways, we attained a heart organoid that incorporates 1) a functional primary coronary vascular plexus, comprising primary/venous CD31^+^/DACH1^+^/NR2F2^+^ and artery-like CD31^+^/DACH1^+^/WT1^+^ ECs, lined by NG2^+^ pericytes and LAM^+^/FIBRO^+^ ECM proteins, which support the maturation of the vascular plexus; and 2) functional SNs (TH^+^/PRPH^+^) that are capable of controlling heart organoid function.

Lineage tracing studies in mouse model have given important insights and established a consensus idea that the primary coronary plexus is mainly derived from both the endocardium and the sinus venosus, and epicardial-derived pericytes are required for coronary vasculature maturation, integrity, growth, and vascular guidance ^23^. Although further validation is needed, we hypothesised that the coronary vascular-like network observed in vascularized EMOs may be derived from a NR2F2^+^ sinus venous-like population of cells that are present in these organoids. Also, although the contribution of epicardium to the ECs that comprise the coronary vasculature of the heart is still debatable, it has been demonstrated in mouse and human fetal hearts, that WT1 is expressed in ECs of both arteries and veins at early stages of development ^34,35^. Lineage tracing of pro-epicardium and septum transversum cells also identified WT1^+^/CD31^+^ cells in the coronary vasculature of the developing mouse heart ^36^. In vascularized EMOs we observed that the DACH1^+^/CD31^+^ ECs, present in the sub-epicardial space and within the myocardial region, express WT1, which may validate the data observed in the previously described models.

The positive impact of epicardium on myocardium growth, compaction and maturation in vivo is well described in the literature ^37,38^. Although we had previously reported the presence of a compact myocardial region in EMOs ^15^, the incorporation of vascularization and the promotion of EMT of epicardial progenitor cells into fibroblasts and pericytes, significantly increased the robustness of the compact myocardial region in EMOs, with increased number of proliferative CMs (NKX2.5^+^/KI67^+^) and ECM deposition. In fact, it is well documented in mouse models that coronary ECs support myocardial compaction and heart wall expansion, and impairment of these developmental process leads to congenital heart defects, including ventricular non-compaction cardiomyopathy ^39^.

The control of heart function through autonomic nervous system modulation is a promising therapeutic strategy for several arrhythmogenic cardiomyopathies. As a proof-of-concept, we showed that iEMOs presented here an adequate response when stimulated with nicotine, by promoting an increased frequency of organoid contraction. This observation not only proves the functional interaction between the SNs and the CMs present in the iEMOs, but also shows the suitability of this model to test therapeutic strategies based on SNs activity manipulation/modulation in a human-based setting. This is particularly relevant since it has been proved that drug responses and disease mechanisms between non-human animals and humans are different.

It is well known that altered density and/or function of heart innervation and vascularization are the most prevalent causes of heart failure, arrhythmogenesis, and sudden cardiac death. In fact, apart from CM death and the subsequent electrophysiological and fibrotic remodelling observed after myocardial infarction (MI) events, the level of cardiac sympathetic nerve fibers loss has been considered the most relevant predictor of consequent ventricular arrhythmias ^40,41^. The re-activation of developmental programs for generating these cells has been emphasized as a promising strategy in regenerative medicine. iEMOs are, therefore, a unique model to unravel crucial signals involved in the normal coronary vascular growth and SNs innervation of the ventricle myocardium, which can be further explored to develop new regenerative medicine-based therapeutic avenues to restore innervation and promote re-vascularization in adult heart after ischemic events.

Additionally, the herein described platform to generate innervated vascularized EMOs, is not only a valuable asset to perform developmental biology studies and validate data previously attained in animal models in a human context, but also a unique vehicle for developmental toxicity studies. iEMOs are generated in a stepwise and in-vivo like environment, that recreates the sequential stages of human heart embryogenesis, including 1) mesendoderm formation, 2) cardiac progenitor cell specification, 3) ventricular myocardium differentiation, 4) pro-epicardium/septum transversum organ formation, 4) epicardial layer establishment, 5) myocardium maturation and compaction, 6) coronary vasculature development, and 7) sympathetic innervation of the myocardium. Defects in the formation of the coronary vasculature can compromise the normal progression of myocardium growth and maturation, therefore contributing to congenital heart defects, and disruption of cardiac sympathetic patterning in the embryo, which can lead to postnatal arrhythmia and cardiac failure.

This work presents a multifactorial platform to generate a modular heart organoid that although exploring a self-organization environment, is complemented with exogenously controlled key steps which ensures the robustness of the process. This strategy favours the generation of organoids presenting reduced variability, which will be crucial to ensure reproducibility of the data acquired through this innovative model.

## LIMITATIONS OF THE STUDY

In this study, the characterization of the epicardial region of V- and P-EMOs lacks depth. We clearly showed that this area comprises WT1, NR2F2 and DACH1 positive cells, however the complete identification of the different subpopulations of cells present within this region still needs to be clarified. Although human based data regarding epicardial and epicardial-derived cells is still not completely disclosed, single cell RNA-seq analysis could elucidate on that regard. Additionally, although we proposed a sinus venous-like NR2F2^+^ cell population as the source of the coronary-like CD31^+^ ECs present in EMOs, further analysis needs to be performed to support this claim. Also, we suggest in this study the presence of at least two subtypes of coronary-like vascular cells in EMOs, namely arterial and venous-like ECs, with different locations within the organoid and with different nuclear staining. However, further validation is needed to confirm this hypothesis. Regarding innervated EMOs, we did not perform a full characterization of the SNSs, and therefore other cell types present in these spheroids, apart from the SNs, can be also playing a role in the assembloid model.

## DECLARATION OF INTERESTS

This paper forms the basis of a patent application (PPP 2024200621851) on which M.A.B. and M.M.D are named as inventors.

## METHODS

### Cell Maintenance

hiPSCs (cell line iPS-DF6-9-9T.B) was maintained in mTeSR^TM^1 (StemCell Technologies) in six-well plates coated with Matrigel™ (Corning). The medium was changed daily. Cells were routinely passaged every three to four days using 0.5 mM EDTA solution (Thermo Fisher Scientific).

### Generation of EMOs

The generation of EMOs starts with the combination of hiPSC-myocardium organoids (MOs) and hiPSC-pro-epicardium organoids (PEO), previously described by our group ^15,42^. Briefly, the differentiation of hiPSC-spheroids (4000 cells/spheroid) was initiated on day 0 (D0) by replacing the mTeSR^TM^1 by RPMI 1640 medium (Thermo Fisher Scientific) with 2%(v/v) B-27 minus insulin (Thermo Fisher Scientific) (RB27-) supplemented with 11 μM CHIR99021 (CHIR). After 24 h, the medium was changed and on D3, cells were cultured in RB27-supplemented with IWP-4 at a final concentration of 5 μM, for two days. At D5, in the case of MOs differentiation, the medium was changed to RB27-, and in the case of PEOs differentiation, 3 μM CHIR, 25 ng/mL BMP4, and 4 μM RA were added from D5 to D7, using DMEM/F12 + 2.5 mM Glutamax + 100 μg/mL of ascorbic acid (DMEM/F12) as basal medium. At D7, aggregates were flushed from the AggreWell™800 plate (StemCell) and transferred to 6-Well ultra-Low Attachment (ULA) plates. Thereafter, the medium was changed every two days until cell harvest. In the case of MOs, the medium used was RPMI 1640 medium supplemented with 2%(v/v) B-27 and in the case of PEOs it was used DMEM/F12 medium.

To establish the co-culture system, D11 MOs and D11 PEOs were dissociated to single cells using 0.25% (v/v) Trypsin-EDTA for 7 min at 37 °C. After cell counting, both cell types were combined at a proportion of 90%MOs:10%PEOs cells and reaggregated in ultra-low 96-well microwell plates. Organoids were cultured in DMEM/F12 medium for six days. After that time, the culture medium was supplemented with VEGFA (50 ng/mL) and PDGFBB (5 ng/mL) factors, alone or combined, from D6 to D10 of co-culture, to evaluate the sole and combined effect of these factors to promote epithelial to mesenchymal transition (EMT) of epicardial progenitor cells and coronary vascularization.

### Differentiation of sympathetic neuron spheroids (SNSs) from hiPSCs

To initiate differentiation of SNSs, hiPSCs were harvested with accutase (Sigma) for 7 min at 37 °C. After dissociation, spheroids of hiPSCs were generated using microwell plates (AggreWell™800, StemCell Technologies) according to the manufacturer’s instructions. Cells were plated at a cell density of 0.3 × 10^6^ cells/well in mTeSR™1 supplemented with 10 μM ROCKi. 24 h later, the differentiation process (based on ^11^) was initiated. Briefly, from D0 to D11 we used 1) E6 basal media from D0 to D3, 2) mixed E6 and N2B27 (Neurobasal™ Plus Medium + B27 Plus + N2 supplement) at (75:25)%, respectively, from D3 to D6 3), and mixed E6 and N2B27 at (50:50)%, from D6 to D8, and 4) only N2B27 from D8-D11. At D0, basal medium was supplemented with 500 nM LDN193189 (LDN) and 10 µM SB431542 (SB) and at D2, the medium was supplemented with 500 nM LDN + 10 µM SB + 3 µM CHIR + 10 µM DAPT. At D3, the medium was supplemented with 10 µM SB + 3 µM CHIR + 10 µM DAPT + 1 µM SAG and aggregates were transferred to ULA 6-well plates. From D4 to D7, the medium was supplemented every day with 3 µM CHIR + 10 µM DAPT + 1 µM SAG. From D7 to D11, the medium was supplemented every day with only 1 µM SAG. At D11, SN progenitor spheroids were singularized using accutase for 10 min at 37 °C. After cell counting, cells were resuspended in N2 maturation media (Neurobasal™ Plus medium + 2% (v/v) B27Plus + 1% (v/v) Glutamax + 1% (v/v) N2 supplement + 0.2 mM Ascorbic Acid + 10 ng/mL NGF + 10 ng/mL BDNF + 10 ng/mL GDNF) supplemented with 10 μM of ROCKi (100 µL/well), and then seeded on a U-shaped ULA 96-well plate. From this point on, the medium was changed every three-four days and maturation was prolonged for 30-40 days.

### Generation of innervated heart assembloids

Heart-innervated assembloids were generated by combining EMOs, obtained after 10 days of co-culture, with SNSs obtained after 30-40 days of maturation in U-shaped ULA 96-well plate (1:1 SNSs/EMOs). These assembloids were cultured in 50:50 N2 maturation medium and DMEM/F12 for 10 days and the medium was changed every 3 days.

### Viral labeling of neural spheroids

To follow SN projections towards the EMO region of the assembloids, day 30 SNSs were labeled with pAAV-CAG-tdTomato virus (Addgene, cat. no. 59462-AAV1). For labeling SNSs, each spheroid was cultured in 40 µL of N2 maturation medium supplemented with 0.5 µL of virus solution in a 96-well plate, overnight. On the next day, 100 µL of N2 maturation medium was added to each well. After 24 h, the SNSs were washed twice with PBS and cultured in fresh N2 maturation medium. Fluorescently labeled SNSs were usually observed seven days after virus infection.

### Immunostaining analysis of EMOs and SNSs

EMOs and SNSs were fixed in 4% PFA in PBS at 4 °C overnight. After PFA removal, cells were stored in PBS at 4 °C for further analysis.

*Slices.* EMOs were incubated in 15%(w/v) sucrose in PBS, at 4 °C overnight. Afterwards, the 3D models were embedded in 7.5%(w/v)/15%(w/v) gelatin/sucrose and frozen in isopentane at −80 °C. Organoid/spheroid sections of 10-12 μm of thickness were cut on a cryostat-microtome (Leica CM3050S, Leica Microsystems), collected on Superfrost™ Microscope Slides and stored at −20 °C. Sections were then de-gelatinized for 45 min in PBS at 37°C. Organoid/spheroid sections were incubated in 0.1 M Glycine (Millipore) for 10 min at RT, permeabilized with 0.1%(v/v) Triton X-100 (Sigma) in PBS at RT for 10 min and blocked with 10%(v/v) fetal bovine serum (FBS, Gibco) in TBST (20 mM Tris-HCl pH 8.0, 150 mM NaCl and 0.05%(v/v) Tween-20, Sigma), at RT for 30 min. Organoids/spheroids were then incubated with the primary antibody diluted in 10%(v/v) FBS in TBST solution (**Table S1**) at 4 °C overnight. Secondary antibodies were added for 30 min in dark, and nuclear counterstaining was performed using 4,6-diamidino-2-phenylindole (DAPI, 1.5 μg/mL, Sigma), at RT for 5 min.

*Whole mount.* EMOs were incubated in 0.1 M Glycine for 30 min at RT, permeabilized with 0.1%(v/v) Triton X-100 (Sigma) in PBS at RT for 30 min and blocked with 10%(v/v) FBS in TBST, at RT for 1 hour. Organoids were then incubated with the primary antibody diluted in 10%(v/v) FBS in TBST solution (**Table S1**) at 4 °C overnight, followed by secondary antibody incubation at 4 °C overnight, in the dark.

### Light sheet Microscopy of vascularized and innervated EMOs

Fluorescence images of vascularized and innervated EMOs were acquired using a Light Sheet Fluorescence Microscope (LSFM), the Zeiss Lightsheet Z.1, with solid states laser lines 488 nm and 561 nm selected for excitation of Alexa-488 and tdTomato or Alexa-546 and associated filter sets 505-545 nm and 575-615 nm, respectively, equipped with a W Plan-Apochromat 20x/1.0 Corr DIC M27 75 mm objective and a pco.edge 5.5 sCMOS camera. The organoids were embedded in 1 %(m/v) low melting-point agarose, and a glass capillary was used as a sample embedding container in the water chamber. Z-stacks from 4 viewing angles were acquired and fused with ZEN (Zeiss) software to generate a 3D volume rendering image of the entire sample.

### Multielectrode Array (MEA) recording of SNSs

SNSs were plated in a Matrigel-coated high-density MEA system comprising 26400 electrodes per well (MaxWell Biosystems AG; MX2-S-6W). SNSs were cultured for 12-20 days before data recording. Network and axon tracking analysis was performed to attain information regarding axon length, neuron conduction velocity and neuron firing rate, using a MaxWell Biosystems software. During recordings, the temperature was kept at 37°C.

### RT-qPCR analysis of SN progenitors

Total RNA was isolated from SN progenitors both at days 6 and 12 of differentiation, using a High Pure RNA Isolation Kit (Roche) according to the manufactureŕs instructions. Total RNA was converted into cDNA with High-Capacity cDNA Reverse Transcription Kit (Thermo Fisher Scientific). PCR reactions were performed with Syber Green Master Mix (nzytech; **Table S2**). Reactions were run in triplicate in ViiA7 Real-Time PCR Systems (Applied BioSystems). For each analyzed time point, gene expression was normalized against the expression of the housekeeping gene glyceraldehyde-3-phosohate dehydrogenase (GAPDH) and results were analysed with the QuantStudioTM RT-PCR Software.

### Calcium transient imaging of EMOs

Fluo-4-AM dye (ThermoFisher) was used for calcium imaging of EMOs. For that, Fluo-4-AM dye-loading solution was prepared according to the manufacturer’s instructions and EMOs were incubated with that solution for 30 min at 37 °C. Afterwards, the Fluo-4-AM dye-loading solution was exchanged by DMEM/F12 medium. Before starting image acquisition, EMOs were incubated at 37°C for another 30 min to stabilize. Videos of beating EMOs were taken for a period of 30–60 s with a ZEISS Cell Observer SD confocal microscope. Calcium transient profiles were analyzed using an in-house Python-developed software.

### Statistical analysis and reproducibility

All presented data regarding IF staining, MEA and PCR analysis were representative of at least three biologically independent experiments per condition/marker. All raw data was collected in Microsoft Excel and statistical analysis was performed with GraphPad software. Statistical significance was evaluated with a two-tailed, unpaired Student’s t test (p < 0.05) when appropriate. The data are presented as mean ± SEM and represent a minimum of three independent experiments. Statistical significance was assigned as not significant (ns), p > 0.05; *p ≤ 0.05; **p ≤ 0.005; ***p ≤ 0.0005. All micrograph images are representative of at least four independent experiments per condition/marker.

## SUPPLEMENTARY FIGURES LEGENDS

**Figure S1:**
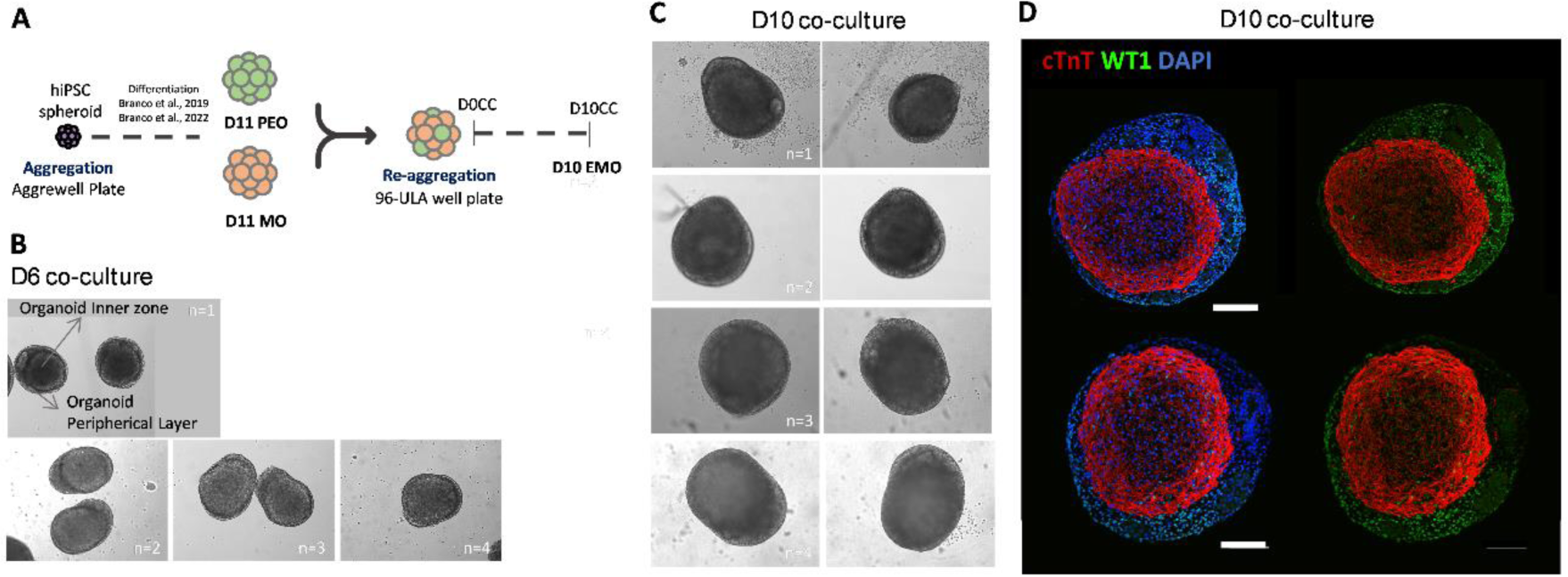
Characterization of control EMOs. **(A)** Scheme illustrating EMOs generation from hiPSC-derived myocardium and pro-epicardium organoids (MO and PEO). **(B-C)** BF images of EMOs after 6 (B) and 10 (C) days of co-culture, for 4 independent assembling experiments, highlighting the reproducibility of the structure of these assembloids. **(D)** Representative IF images of EMOs after 10 days of co-culture, highlighting the two main regions identified in these organoids, the epicardium layer (WT1^+^) and the myocardium zone (cTnT^+^). Scale bars, 100 µm. Related with Figure 1.

**Figure S2:**
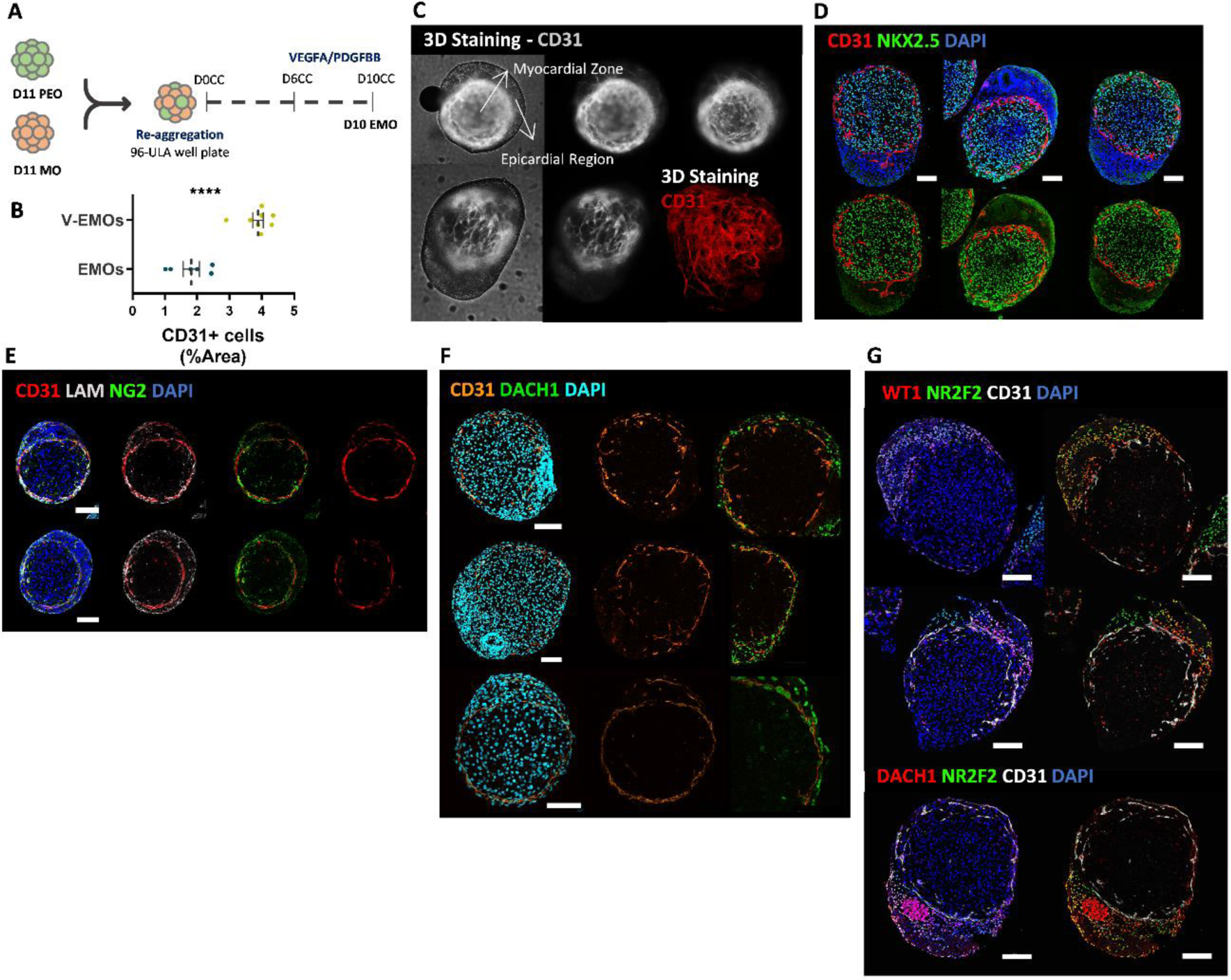
Characterization of EMOs after VEGF signalling modulation. **(A)** Scheme illustrating EMOs generation with VEGFA and PDGFBB stimulation from day 6 to day 10 of co-culture. **(B)** Percentage of EMOs area that stains for the EC marker CD31, with and without VEGFA supplementation. Data was obtained from representative IF images of at least 6 independent experiments. **(C)** Representative IF staining of VEGFA-treated EMOs (3D and organoid slices), highlighting the development of a CD31^+^ network of vascular cells surrounding the myocardium region. **(D-G)** Representative IF images of V-EMOs (organoid slices), highlighting the sprouting of CD31^+^ cells towards the myocardium NKX2.5^+^ region (D), the co-staining of CD31 network with the ECM molecule laminin and the pericyte marker NG2 (E), and the staining for DACH1, WT1 and NR2F2 in CD31^+^ cells and within the epicardium region (F and G). Related with Figure 1.

**Figure S3:**
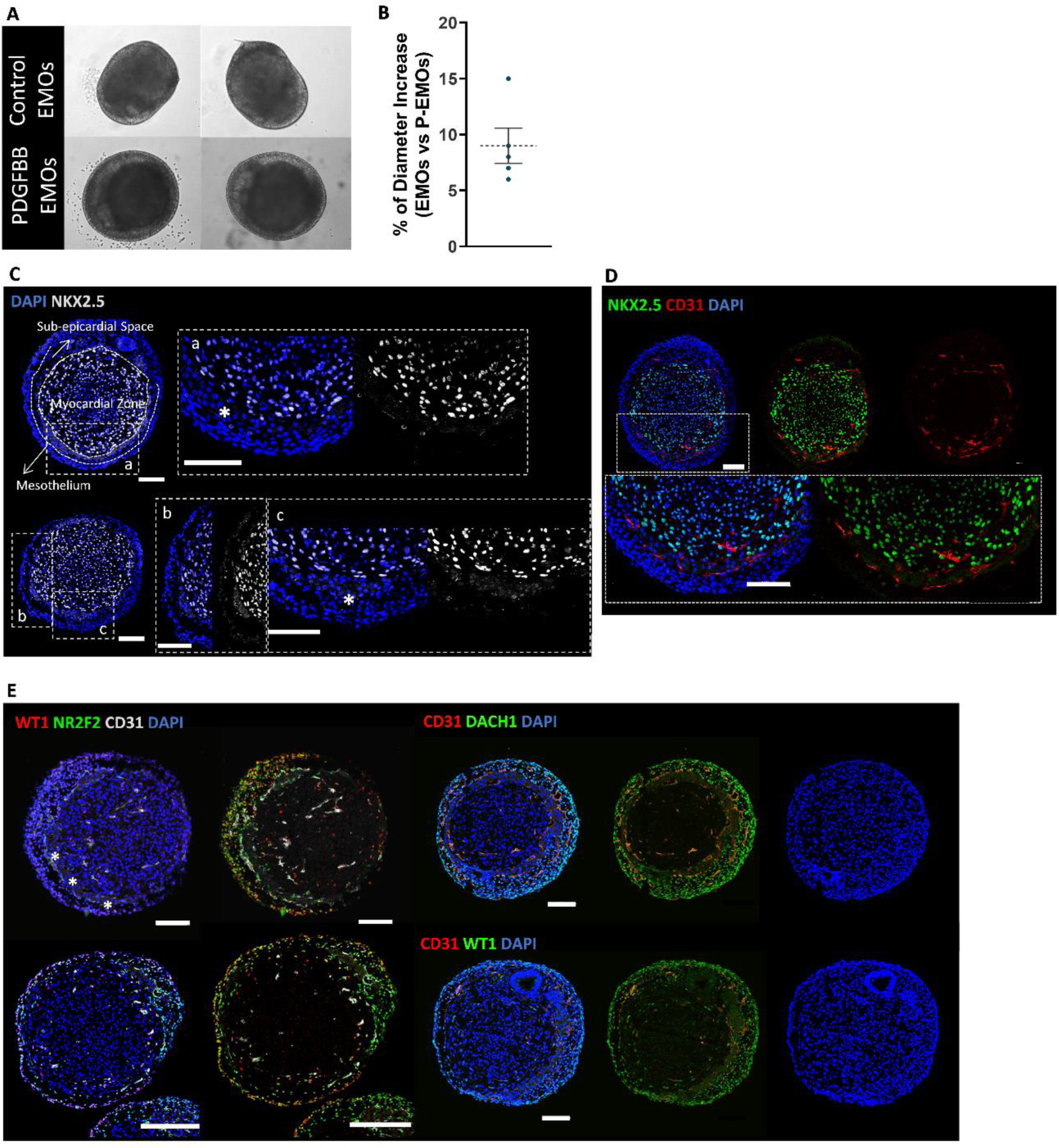
Characterization of EMOs after PDGF-β signalling modulation. **(A)** Representative BF images of control and P-EMOs after 10 days of culture. **(B)** Quantification of P-EMO size. Data was obtained from representative BF images of 5 independent experiments. **(C-D)** Representative IF images of the epicardium region in P-EMOs, (organoid slices), highlighting the generation of two distinct layers in the epicardium region, the mesothelium (WT1^+^/NR2F2^+^/DACH1^+^ cells) and the sub-epicardial space (WT1^−/+low^, NR2F2^+^ and DACH1^+^ cells). **(E)** Representative IF images of P-EMOs (organoid slices), highlighting the localization of the CD31^+^/WT1^+^ ECs within the sub-epicardium space and the compact myocardium region. Scale bars, 100 µm. Related with Figure 1.

**Figure S4:**
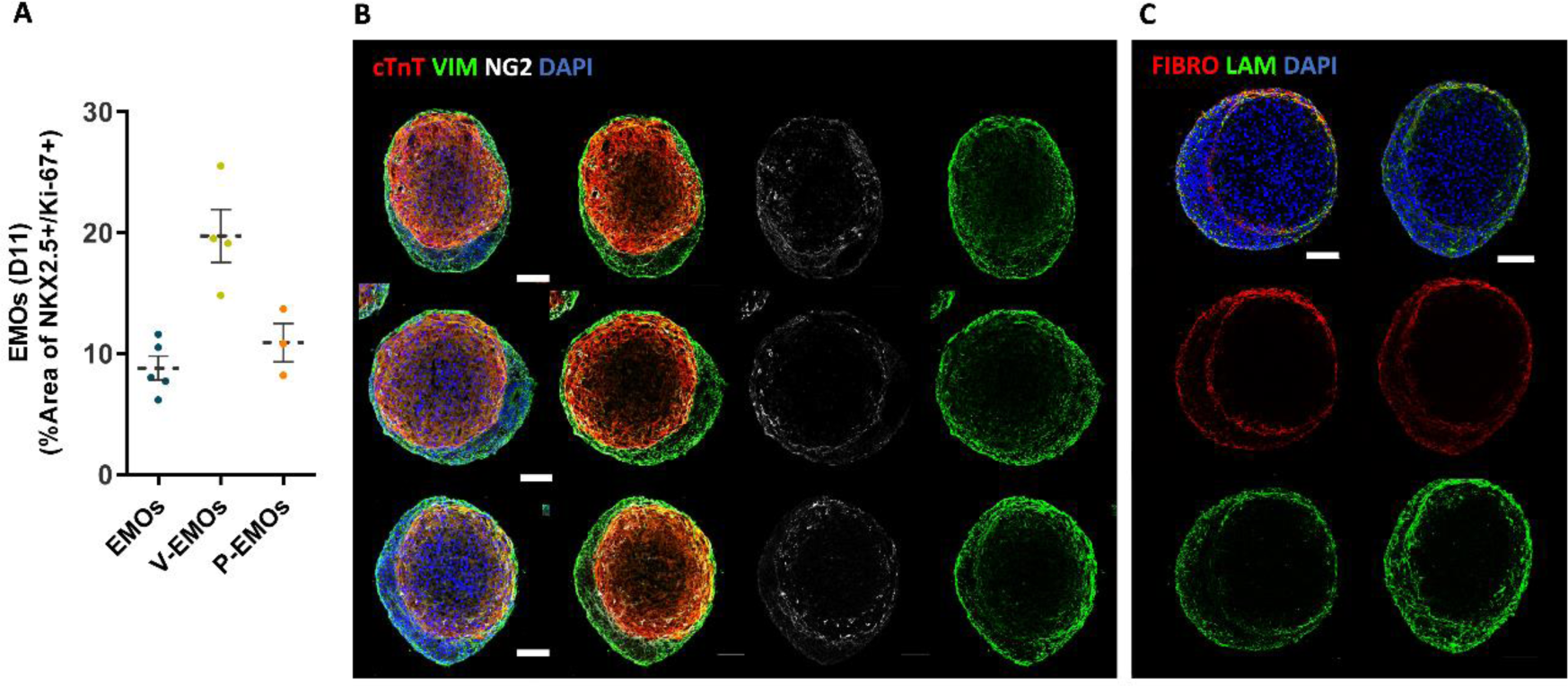
Characterization of compact myocardium in V- and P-EMOs. **(A)** Quantification of proliferative CMs within the myocardial region in EMOs, and V- and P-EMOs. Data was attained from representative IF images of at least 3 independent experiments. **(B-C)** Representative IF images of V-EMOs, (organoid slices), highlighting the compact myocardial region, and the co-localization with ECM (LAM, FIBRO), pericytes (NG2) and fibroblast (VIM) markers. Scale bars, 100 µm. Related with Figure 2.

**Figure S5:**
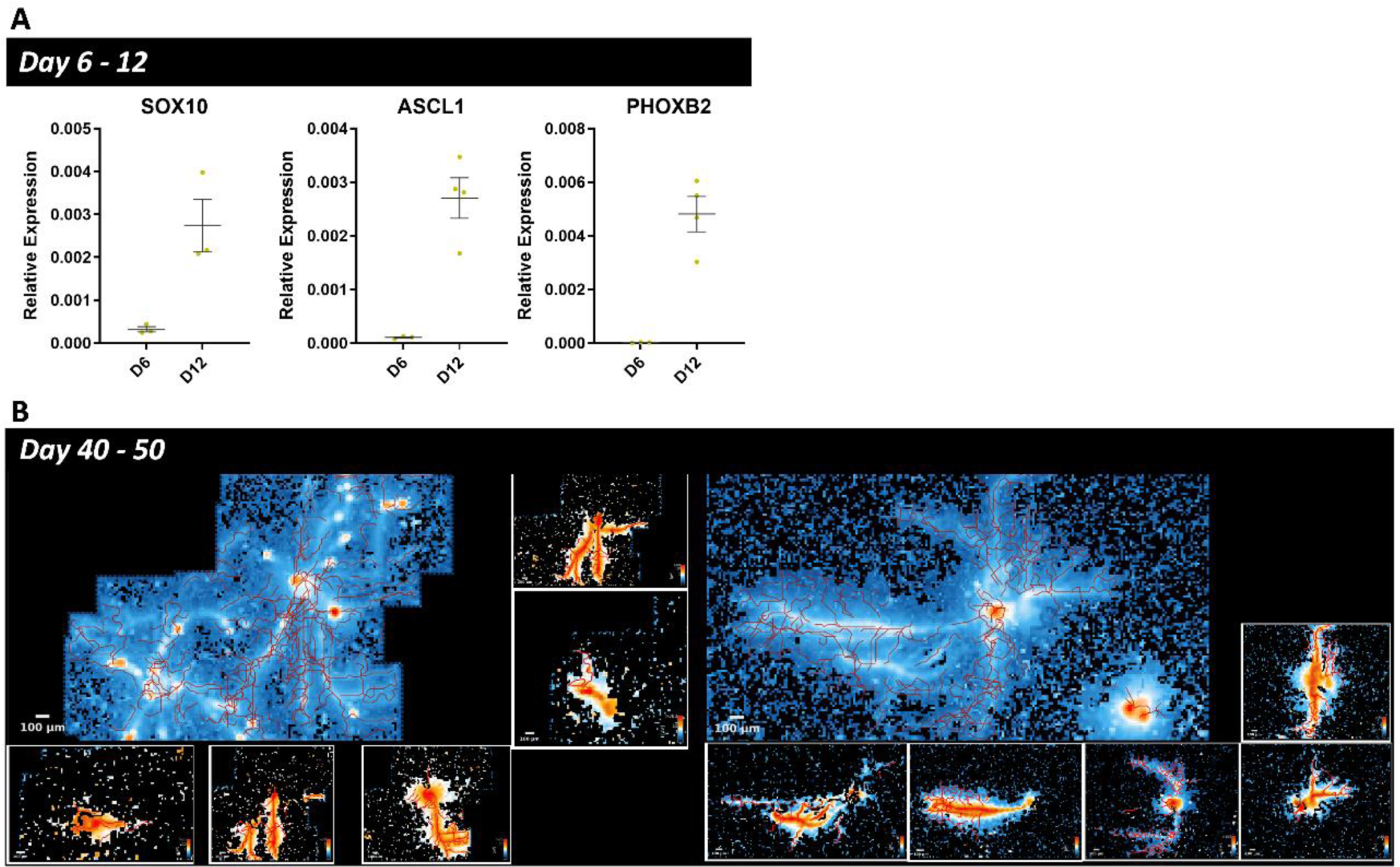
Molecular and functional characterization of SNSs. **(A)** Expression profile of SOX10, ASCL1 and PHOXB2 genes at days 6 and 12 of SNSs differentiation. Values are normalized to GAPDH. Data are represented as mean ± SEM of n = 4 independent experiments. **(B)** Representative neuronal network maps obtained from MEA analysis of SNSs after 50 days of differentiation off chip plus 8-12 days of differentiation on chip, highlighting the identified neuronal branches. Related with Figure 3

## SUPPLEMENTARY INFORMATION TITLES

Table S1 – List of antibodies used in Immunostaining analysis

Table S2 – List of primers used in real time PCR analysis

Video S1 – Coronary vascularization of V-EMOs, related to Figure 1

## ACKNOWLEDGMENTS

We thank Instituto de Medicina Molecular (iMM) for the access to the bioimaging facility. We acknowledge funding received from the national foundation FCT - Fundação para a Ciência e a Tecnologia, Portugal, through Institute for Bioengineering and Biosciences - iBB (UIDB/04565/2020 and UIDP/04565/2020), through Associate Laboratory Institute for Health and Bioeconomy – i4HB (LA/P/0140/2020), and through the project 2022.02251.PTDC granted to M.A.B. We also acknowledge funding received from Agência Nacional de Inovação and European Union (REC05-i02 –N° 01/C05-i02/2022 and 01/C05-i02/2022).

## Notes

### Competing Interest Statement

The authors have declared no competing interest.

## REFERENCES

1. Giacomelli, E., Meraviglia, V., Campostrini, G., Cochrane, A., Cao, X., van Helden, R.W.J., Krotenberg Garcia, A., Mircea, M., Kostidis, S., Davis, R.P., et al. (2020). Human-iPSC-Derived Cardiac Stromal Cells Enhance Maturation in 3D Cardiac Microtissues and Reveal Non-cardiomyocyte Contributions to Heart Disease. Cell Stem Cell 26, 862–879.e11. 10.1016/j.stem.2020.05.004.

2. Arslan, U., Brescia, M., Meraviglia, V., Nahon, D.M., van Helden, R.W.J., Stein, J.M., van den Hil, F.E., van Meer, B.J., Vila Cuenca, M., Mummery, C.L., et al. (2023). Vascularized hiPSC-derived 3D cardiac microtissue on chip. Stem Cell Reports 18, 1394–1404. 10.1016/j.stemcr.2023.06.001.

3. Yang, J., Lei, W., Xiao, Y., Tan, S., Yang, J., Lin, Y., Yang, Z., Zhao, D., Zhang, C., Shen, Z., et al. (2024). Generation of human vascularized and chambered cardiac organoids for cardiac disease modelling and drug evaluation. Cell Prolif., 1–13. 10.1111/cpr.13631.

4. Voges, H.K., Foster, S.R., Reynolds, L., Parker, B.L., Devilée, L., Quaife-Ryan, G.A., Fortuna, P.R.J., Mathieson, E., Fitzsimmons, R., Lor, M., et al. (2023). Vascular cells improve functionality of human cardiac organoids. Cell Rep. 42, 112322. 10.1016/j.celrep.2023.112322.

5. Hernandez, I., Ramirez, S.P., Salazar, W. V., Mendivil, S., Guevara, A., Patel, A., Loyola, C.D., Dorado, Z.N., and Joddar, B. (2023). A Semi-Three-Dimensional Bioprinted Neurocardiac System for Tissue Engineering of a Cardiac Autonomic Nervous System Model. Bioengineering 10, 834. 10.3390/bioengineering10070834.

6. Li, N., Edel, M., Liu, K., Denning, C., Betts, J., Neely, O.C., Li, D., and Paterson, D.J. (2023). Human induced pluripotent stem cell-derived cardiac myocytes and sympathetic neurons in disease modelling. Philos. Trans. R. Soc. B Biol. Sci. 378. 10.1098/rstb.2022.0173.

7. Sakai, K., Shimba, K., Ishizuka, K., Yang, Z., Oiwa, K., Takeuchi, A., Kotani, K., and Jimbo, Y. (2017). Functional innervation of human induced pluripotent stem cell-derived cardiomyocytes by co-culture with sympathetic neurons developed using a microtunnel technique. Biochem. Biophys. Res. Commun. 494, 138–143. 10.1016/j.bbrc.2017.10.065.

8. Takeuchi, A., Nakafutami, S., Tani, H., Mori, M., Takayama, Y., Moriguchi, H., Kotani, K., Miwa, K., Lee, J.K., Noshiro, M., et al. (2011). Device for co-culture of sympathetic neurons and cardiomyocytes using microfabrication. Lab Chip 11, 2268–2275. 10.1039/c0lc00327a.

9. Häkli, M., Jäntti, S., Joki, T., Sukki, L., Tornberg, K., Aalto-Setälä, K., Kallio, P., Pekkanen-Mattila, M., and Narkilahti, S. (2022). Human Neurons Form Axon-Mediated Functional Connections with Human Cardiomyocytes in Compartmentalized Microfluidic Chip. Int. J. Mol. Sci. 23. 10.3390/ijms23063148.

10. Oiwa, K., Shimba, K., Numata, T., Takeuchi, A., Kotani, K., and Jimbo, Y. (2016). A device for co-culturing autonomic neurons and cardiomyocytes using micro-fabrication techniques. Integr. Biol. (United Kingdom) 8, 341–348. 10.1039/c5ib00273g.

11. Winbo, A., Ramanan, S., Eugster, E., Jovinge, S., Skinner, J.R., and Montgomery, J.M. (2020). Functional coculture of sympathetic neurons and cardiomyocytes derived from human-induced pluripotent stem cells. Am. J. Physiol. - Hear. Circ. Physiol. 319, H927–H937. 10.1152/AJPHEART.00546.2020.

12. Takayama, Y., Kushige, H., Akagi, Y., Suzuki, Y., Kumagai, Y., and Kida, Y.S. (2020). Selective Induction of Human Autonomic Neurons Enables Precise Control of Cardiomyocyte Beating. Sci. Rep. 10, 1–13. 10.1038/s41598-020-66303-3.

13. Oh, Y., Cho, G.-S., Li, Z., Hong, I., Zhu, R., Kim, M.-J., Kim, Y.J., Tampakakis, E., Tung, L., Huganir, R., et al. (2016). Functional Coupling with Cardiac Muscle Promotes Maturation of hPSC-Derived Sympathetic Neurons. Cell Stem Cell 19, 95–106. 10.1016/j.stem.2016.05.002.

14. Nam, J., Onitsuka, I., Hatch, J., Uchida, Y., Ray, S., Huang, S., Li, W., Zang, H., Ruiz-Lozano, P., and Mukouyama, Y.S. (2013). Coronary veins determine the pattern of sympathetic innervation in the developing heart. Dev. 140, 1475–1485. 10.1242/dev.087601.

15. Branco, M.A., Dias, T.P., Cabral, J.M.S., Pinto-do-Ó, P., and Diogo, M.M. (2022). Human multilineage pro-epicardium/foregut organoids support the development of an epicardium/myocardium organoid. Nat. Commun. 13, 6981. 10.1038/s41467-022-34730-7.

16. Mellgren, A.M., Smith, C.L., Olsen, G.S., Eskiocak, B., Zhou, B., Kazi, M.N., Ruiz, F.R., Pu, W.T., and Tallquist, M.D. (2008). Platelet-Derived Growth Factor Receptor β Signaling Is Required for Efficient Epicardial Cell Migration and Development of Two Distinct Coronary Vascular Smooth Muscle Cell Populations. Circ. Res. 103, 1393–1401. 10.1161/CIRCRESAHA.108.176768.

17. Smith, C.L., Baek, S.T., Sung, C.Y., and Tallquist, M.D. (2011). Epicardial-Derived Cell Epithelial-to-Mesenchymal Transition and Fate Specification Require PDGF Receptor Signaling. Circ. Res. 108. 10.1161/CIRCRESAHA.110.235531.

18. Iyer, D., Gambardella, L., Bernard, W.G., Serrano, F., Mascetti, V.L., Pedersen, R.A., Talasila, A., and Sinha, S. (2015). Robust derivation of epicardium and its differentiated smooth muscle cell progeny from human pluripotent stem cells. Development 142, 1528–1541. 10.1242/dev.119271.

19. Bonet, F., Pereira, P.N.G., Bover, O., Marques, S., Inácio, J.M., and Belo, J.A. (2018). CCBE1 is required for coronary vessel development and proper coronary artery stem formation in the mouse heart. Dev. Dyn. 247, 1135–1145. 10.1002/dvdy.24670.

20. Chen, H.I., Sharma, B., Akerberg, B.N., Numi, H.J., Kivelä, R., Saharinen, P., Aghajanian, H., McKay, A.S., Bogard, P.E., Chang, A.H., et al. (2014). The sinus venosus contributes to coronary vasculature through VEGFC-stimulated angiogenesis. Development 141, 4500–4512. 10.1242/dev.113639.

21. Chang, A.H., Raftrey, B.C., D’Amato, G., Surya, V.N., Poduri, A., Chen, H.I., Goldstone, A.B., Woo, J., Fuller, G.G., Dunn, A.R., et al. (2017). DACH1 stimulates shear stress-guided endothelial cell migration and coronary artery growth through the CXCL12–CXCR4 signaling axis. Genes Dev. 31, 1308–1324. 10.1101/gad.301549.117.

22. Raftrey, B., Williams, I.M., Rios Coronado, P.E., Fan, X., Chang, A.H., Zhao, M., Roth, R., Trimm, E., Racelis, R., D’Amato, G., et al. (2021). Dach1 Extends Artery Networks and Protects Against Cardiac Injury. Circ. Res. 129, 702–716. 10.1161/CIRCRESAHA.120.318271.

23. Quijada, P., Trembley, M.A., Misra, A., Myers, J.A., Baker, C.D., Pérez-Hernández, M., Myers, J.R., Dirkx, R.A., Cohen, E.D., Delmar, M., et al. (2021). Coordination of endothelial cell positioning and fate specification by the epicardium. Nat. Commun. 12, 4155. 10.1038/s41467-021-24414-z.

24. Zhang, M., Pu, W., Li, J., Han, M., Han, X., Zhang, Z., Lv, Z., Smart, N., Wang, L., and Zhou, B. (2023). Coronary vessels contribute to de novo endocardial cells in the endocardium-depleted heart. Cell Discov. 9, 4. 10.1038/s41421-022-00486-z.

25. Pereira, F.A., Qiu, Y., Zhou, G., Tsai, M.-J., and Tsai, S.Y. (1999). The orphan nuclear receptor COUP-TFII is required for angiogenesis and heart development. Genes Dev. 13, 1037–1049. 10.1101/gad.13.8.1037.

26. Volz, K.S., Jacobs, A.H., Chen, H.I., Poduri, A., McKay, A.S., Riordan, D.P., Kofler, N., Kitajewski, J., Weissman, I., and Red-Horse, K. (2015). Pericytes are progenitors for coronary artery smooth muscle. Elife 4, 1–22. 10.7554/eLife.10036.

27. Tian, X., Hu, T., Zhang, H., He, L., Huang, X., Liu, Q., Yu, W., He, L., Yang, Z., Zhang, Z., et al. (2013). Subepicardial endothelial cells invade the embryonic ventricle wall to form coronary arteries. Cell Res. 23, 1075–1090. 10.1038/cr.2013.83.

28. Drakhlis, L., Biswanath, S., Farr, C.-M., Lupanow, V., Teske, J., Ritzenhoff, K., Franke, A., Manstein, F., Bolesani, E., Kempf, H., et al. (2021). Human heart-forming organoids recapitulate early heart and foregut development. Nat. Biotechnol. 39, 737–746. 10.1038/s41587-021-00815-9.

29. Lewis-Israeli, Y.R., Wasserman, A.H., Gabalski, M.A., Volmert, B.D., Ming, Y., Ball, K.A., Yang, W., Zou, J., Ni, G., Pajares, N., et al. (2021). Self-assembling human heart organoids for the modeling of cardiac development and congenital heart disease. Nat. Commun. 12, 5142. 10.1038/s41467-021-25329-5.

30. Hofbauer, P., Jahnel, S.M., Papai, N., Giesshammer, M., Deyett, A., Schmidt, C., Penc, M., Tavernini, K., Grdseloff, N., Meledeth, C., et al. (2021). Cardioids reveal self-organizing principles of human cardiogenesis. Cell 184, 3299–3317. 10.1016/j.cell.2021.04.034.

31. Volmert, B., Kiselev, A., Juhong, A., Wang, F., Riggs, A., Kostina, A., O’Hern, C., Muniyandi, P., Wasserman, A., Huang, A., et al. (2023). A patterned human primitive heart organoid model generated by pluripotent stem cell self-organization. Nat. Commun. 14, 8245. 10.1038/s41467-023-43999-1.

32. Schmidt, C., Deyett, A., Ilmer, T., Haendeler, S., Torres Caballero, A., Novatchkova, M., Netzer, M.A., Ceci Ginistrelli, L., Mancheno Juncosa, E., Bhattacharya, T., et al. (2023). Multi-chamber cardioids unravel human heart development and cardiac defects. Cell 186, 5587–5605.e27. 10.1016/j.cell.2023.10.030.

33. Ge, Y., Smits, A.M., van Munsteren, J.C., Gittenberger-de Groot, A.C., Poelmann, R.E., van Brakel, T.J., Schalij, M.J., Goumans, M.-J., DeRuiter, M.C., and Jongbloed, M.R.M. (2020). Human epicardium-derived cells reinforce cardiac sympathetic innervation. J. Mol. Cell. Cardiol. 143, 26–37. 10.1016/j.yjmcc.2020.04.006.

34. Duim, S.N., Smits, A.M., Kruithof, B.P.T., and Goumans, M.J. (2016). The roadmap of WT1 protein expression in the human fetal heart. J. Mol. Cell. Cardiol. 90, 139–145. 10.1016/j.yjmcc.2015.12.008.

35. Duim, S.N., Kurakula, K., Goumans, M.J., and Kruithof, B.P.T. (2015). Cardiac endothelial cells express Wilms’ tumor-1. Wt1 expression in the developing, adult and infarcted heart. J. Mol. Cell. Cardiol. 81, 127–135. 10.1016/j.yjmcc.2015.02.007.

36. Cano, E., Carmona, R., Ruiz-Villalba, A., Rojas, A., Chau, Y.Y., Wagner, K.D., Wagner, N., Hastie, N.D., Muñoz-Chápuli, R., and Pérez-Pomares, J.M. (2016). Extracardiac septum transversum/proepicardial endothelial cells pattern embryonic coronary arterio-venous connections. Proc. Natl. Acad. Sci. U. S. A. 113, 656–661. 10.1073/pnas.1509834113.

37. Li, P., Cavallero, S., Gu, Y., Chen, T.H.P., Hughes, J., Hassan, A.B., Brüning, J.C., Pashmforoush, M., and Sucov, H.M. (2011). IGF signaling directs ventricular cardiomyocyte proliferation during embryonic heart development. Development 138, 1795–1805. 10.1242/dev.054338.

38. Brade, T., Kumar, S., Cunningham, T.J., Chatzi, C., Zhao, X., Cavallero, S., Li, P., Sucov, H.M., Ruiz-Lozano, P., and Duester, G. (2011). Retinoic acid stimulates myocardial expansion by induction of hepatic erythropoietin which activates epicardial Igf2. Development 138, 139–148. 10.1242/dev.054239.

39. Rhee, S., Chung, J.I., King, D.A., D’amato, G., Paik, D.T., Duan, A., Chang, A., Nagelberg, D., Sharma, B., Jeong, Y., et al. (2018). Endothelial deletion of Ino80 disrupts coronary angiogenesis and causes congenital heart disease. Nat. Commun. 9, 368. 10.1038/s41467-017-02796-3.

40. Tapa, S., Wang, L., Francis Stuart, S.D., Wang, Z., Jiang, Y., Habecker, B.A., and Ripplinger, C.M. (2020). Adrenergic supersensitivity and impaired neural control of cardiac electrophysiology following regional cardiac sympathetic nerve loss. Sci. Rep. 10, 18801. 10.1038/s41598-020-75903-y.

41. Gardner, R.T., Wang, L., Lang, B.T., Cregg, J.M., Dunbar, C.L., Woodward, W.R., Silver, J., Ripplinger, C.M., and Habecker, B.A. (2015). Targeting protein tyrosine phosphatase σ after myocardial infarction restores cardiac sympathetic innervation and prevents arrhythmias. Nat. Commun. 6, 6235. 10.1038/ncomms7235.

42. Branco, M.A., Cotovio, J.P., Rodrigues, C.A.V., Vaz, S.H., Fernandes, T.G., Moreira, L.M., Cabral, J.M.S., and Diogo, M.M. (2019). Transcriptomic analysis of 3D Cardiac Differentiation of Human Induced Pluripotent Stem Cells Reveals Faster Cardiomyocyte Maturation Compared to 2D Culture. Sci. Rep. 9, 1–13. 10.1038/s41598-019-45047-9.

