## Supplementary figures and images for "Recreating coronary vascularization and sympathetic innervation of myocardium on a human pluripotent stem cell-derived heart organoid"

### Table S1

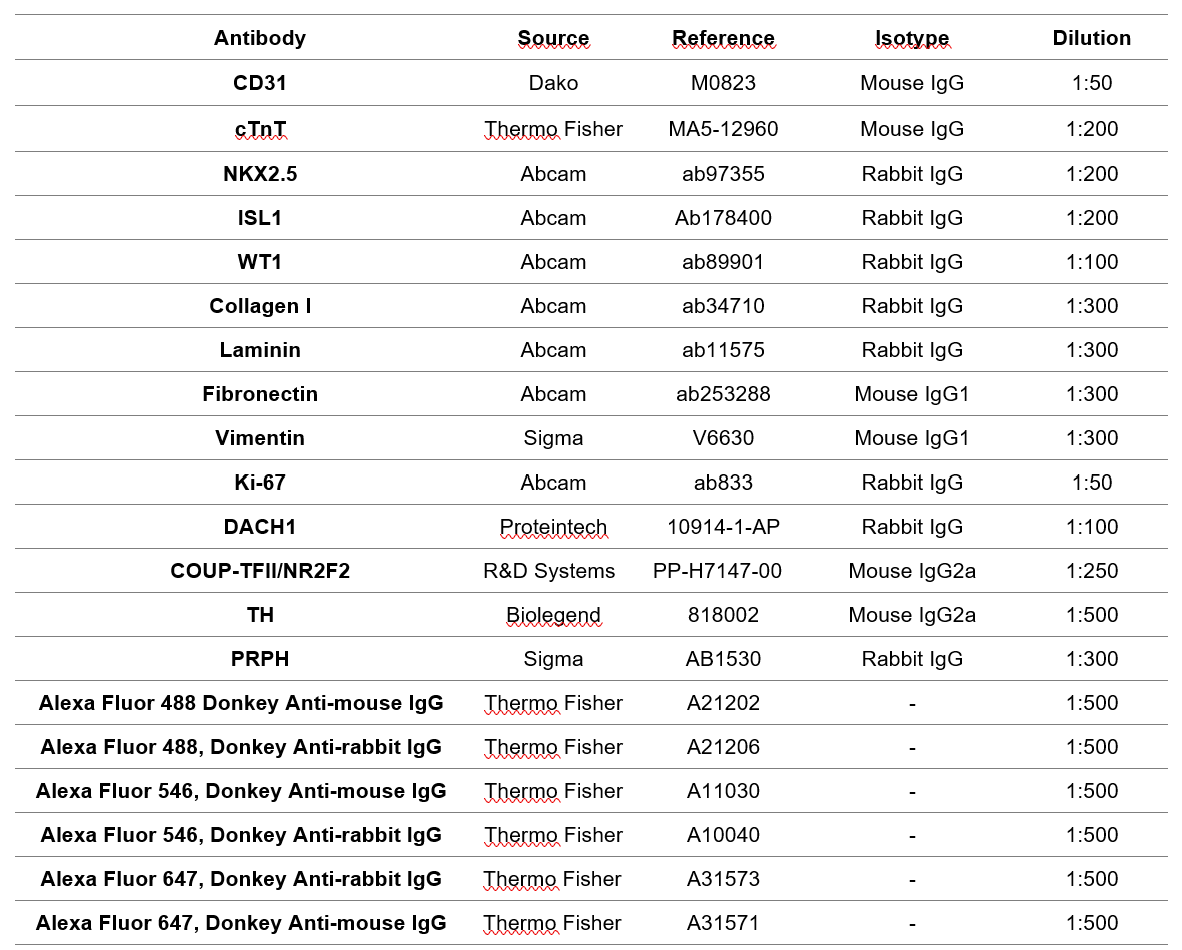

### Table S2

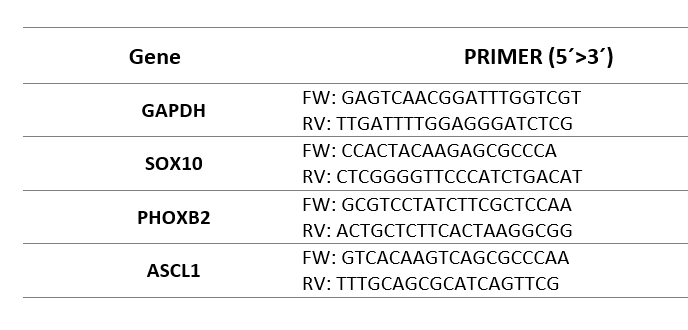
